# Development of transient expression assay for *Cannabis sativa* which revealed differential *Agrobacterium* susceptibility among cannabis cultivars

**DOI:** 10.1101/2020.03.12.988618

**Authors:** Alexei Sorokin, Narendra Singh Yadav, Daniel Gaudet, Igor Kovalchuk

## Abstract

In plant biology, transient expression analysis plays a vital role to provide a fast method to study the gene of interest and subsequently leads the path to develop an improved crop variety with better agronomic traits. In this study, we have reported a rapid and efficient method for transient expression in *Cannabis sativa* seedlings using *Agrobacterium tumefaciens*-mediated transformation. *A. tumefaciens* strain EHA105 carrying the pCAMBIA1301 construct with *uid*A gene was used to transform cannabis seedlings and the GUS assay was used to detect the *uid*A expression. A 1% hydrogen peroxide (H_2_O_2_) solution was used for both seed sterilization and rapid germination steps. Transient transformation revealed that both cotyledons and young true leaves are amenable to transformation. Comparison to *Nicotiana tabacum* (tobacco) showed that cannabis seedlings were less susceptible to transformation with *Agrobacterium tumefaciens*. The susceptibility to *Agrobacterium* infection also varied with the different cannabis cultivars. The method established in this study has potential to be an important tool for gene-function studies and genetic improvement in cannabis.

## Introduction

*Cannabis sativa* is an annual dioecious herb that belongs to family *Cannabaceae*. Historically, cannabis has been widely cultivated as a source of seed oil, fiber and intoxicating resin. First written evidence of using cannabis in medicinal practices is described in the compendium of Chinese medicinal herbs by Emperor Shen Nung, dated 2737 B.C.E. [1]. In the last decades, the therapeutic potential of cannabinoids has been reported for the treatment of a range of human diseases from complex neurological diseases to cancer [2]. Although cannabis is best known for the psychoactive compound D9-tetrahydrocannabinol (THC), it also contains varying levels of non-psychoactive cannabinoids such as cannabidiol (CBD), cannabigerol (CBG), D9-tetrahydrocannabivarin (THCV), and cannabichromene (CBC), that show promising therapeutic properties and in some cases mitigate the psychoactive effects of THC [3].

Considering the enormous economic importance, it is worthy to study the functional genomics of cannabis. The transient expression analysis is an important tool for functional genomics study. *Agrobacterium*-mediated transformation is commonly used to achieve both transient and stable gene expression in plants. Wahby et al. [4] reported that *C. sativa* hypocotyl tissues exhibited high susceptibility to *Agrobacterium* infection/transformation than other tissues. Recently, Chaohua et al. [5] established regeneration protocol that uses cotyledons of *C. sativa* as an explant. Considering aforementioned reports, we decided to use intact cannabis seedlings for establishment of transient expression protocol. Such protocol can be used for functional genomics study and for the development of stable transformation protocol. In this study, we have reported an efficient and reproducible method for transient expression analysis in *C. sativa* seedlings using *Agrobacterium tumefaciens*-mediated transformation and demonstrated that cannabis is less susceptible to *Agrobacterium* transformation than tobacco. Further, we also displayed that susceptibility to *Agrobacterium* infection also varied with the different cannabis cultivars.

## Materials and Methods

### Materials

#### A. Biological materials

1. *Agrobacterium tumefaciens* strain (EHA105) carrying binary vector pCAMBIA1301 with uidA gene.
2. *Cannabis sativa* (Candida CD-1, Night in Gale, Green Crack CBD, and Holy Grail × CD-1 cultivars) and *Nicotiana benthamiana* seeds.

#### B. Chemicals

1. Hydrogen Peroxide 30 % (Merck®, catalog number: 1072091000)
2. Agrobacterium liquid growth medium (YEP liquid medium) (see Recipes)
3. Agrobacterium liquid induction medium (see Recipes)
4. Histochemical GUS staining solution (see Recipes)
5. MS solid media (see Recipes)
6. MgSO4 (Sigma-Aldrich, catalog number: MX0075-1)
7. Acetosyringone (Sigma-Aldrich, catalog number: D134406)
8. Murashige & Skoog Basal Medium with Vitamins (PhytoTechnology Laboratories®, catalog number: M519)
9. Kanamycin sulfate (PhytoTechnology Laboratories®, catalog number: K378)
10. Rifampicin (Sigma-Aldrich, catalog number: R3501)
11. Selective antibiotics: Kanamycin, Rifampicin
12. 70% Ethanol
13. Sucrose (Sigma-Aldrich, catalog number: S0389)
14. MES (Sigma-Aldrich, catalog number: M3671)
15. Agar
16. Yeast extract
17. NaCl
18. Peptone
19. EDTA (pH 8.0) (Sigma-Aldrich, catalog number: E9884)
20. Sodium phosphate buffer (pH 7.0)
21. Triton X-100 (Sigma-Aldrich, catalog number: 234729)
22. Potassium ferricyanide (Sigma-Aldrich, catalog number: 702587)
23. Potassium ferrocyanide (Sigma-Aldrich, catalog number: P3289)
24. X-Gluc (Sigma-Aldrich, catalog number: R0852)

#### C. Plasticware

1. Sterile empty 100 × 15 mm Petri plates (VWR International, catalog number: 25384-342)
2. Sterile disposable 50 ml screw-cap centrifuge tubes (BD, Falcon^™^, catalog number: 352070)
3. Plastic pipette tips (20, 200, and 1,000 μl)
4. Disposable Cuvettes
5. Sterile filter papers

#### D. Equipment

1. Spectrophotometer
2. Allegra Benchtop Centrifuge X-12 (Beckman Coulter)
3. Micro-centrifuge
4. Laminar flow hood
5. Eppendorf Research^®^ plus 10, 20, 200, and 1,000 μl
6. Analytical balance
7. Top loading electronic balance
8. pH meter
9. Vortex mixer
10. Freezer (−80 °C) (e.g. New Brunswick, model:)
11. Sterile forceps and scalpel (sterilized by heat treatment using a Bunsen burner)
12. Sterile inoculating loop
13. A desiccator attached to a vacuum pump (Brinkman DistiVac)
14. 
15. Growth chamber
16. Shaker incubator (28 °C, 220 rpm)
17. Incubator 37°C
18. Fluorescent microscope (Zeiss Observer Z1)

### Methods

#### A. Germination and seedlings development (performed under sterile conditions)

1. For germination, seeds were soaked in a 1% hydrogen peroxide solution incubated overnight for 24 hrs at room temperature in the dark. The following day, radicles with hypocotyl are visible (Figure 1).
2. Transfer germinated seeds into fresh 1% H_2_O_2_ solution and further incubate for 3-4 days until cotyledons have fully opened and two early true leaves are visible.
3. Remove remaining seed coats using sterile scalpel and forceps.
4. Sterilize seedlings without seed coats by soaking them in 1% hydrogen peroxide for 5 min.
5. Prior to transformation rinse seedlings in sterile water 3 times to remove remaining hydrogen peroxide.

**Figure 1:**
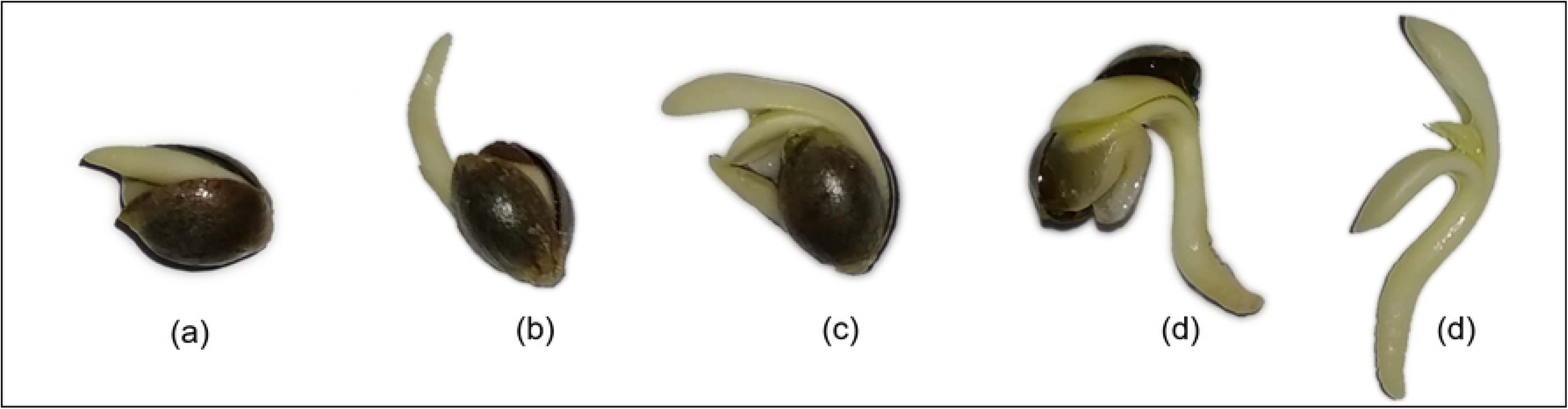
Various germination development stages of cannabis seedlings in 1% hydrogen peroxide solution. (a) 24 hours, cannabis embryo absorbs water until radicle breaks through the seed coat. (b) 36 hours, radicle further development and appearance of hypocotyl. (c) 48 hours, cotyledons emerge. (d) 60 hours, seedling almost leave the seed coat. (e) 72 hours, two fully opened cotyledons and two early true leaves are visible.

#### B. Preparation of Agrobacterium cells culture (all steps performed under sterile conditions)

1. Two days before transformation, inoculate 100 ml of YEP (containing 50 μg/mL Kanamycin and 25 μg/mL Rifampicin) with *Agrobacterium* from glycerol stock and culture at 28°C in an incubator shaker 220 rpm overnight.
2. Next day centrifuge the *Agrobacterium* cells culture at 4,000 × g for 15 min at RT.
3. Remove supernatant and add 3 ml of 10 mM MgSO4, resuspend the *Agrobacterium* pellet.
4. Repeat steps 2 and 3.
5. Centrifuge a third time, remove supernatant.
6. Resuspend the *Agrobacterium* pellet in an appropriate volume of induction medium (MS liquid media) so that the final OD600=0.6.
7. Add 100 mM acetosyringone to final concentration 100 μM.

#### C. Co-cultivation (all steps performed under sterile conditions)

1. Place sterilized seedlings in 50 ml Falcon tubes with 30 ml of the *Agrobacterium* cells suspension *(Agrobacterium* cells in induction medium supplemented with acetosyringone).
2. Place the tubes into a sterile vacuum chamber and apply vacuum for 10-20 min.
3. Transfer seedlings to a sterile filter paper to remove the excess *Agrobacterium* cell culture.
4. Transfer the seedlings to 90 mm petri dishes containing MS media (10 seedlings per plate). Spread them evenly on the plate using forceps. Seal the Petri dishes with parafilm.
5. Co-cultivate the seedlings and the *Agrobacterium* cells for three days in the dark at 25°C.
6. After co-cultivation, seedlings can be used directly for GUS staining or can be frozen at −80°C for further analysis e.g. MUG assay, PCR analysis.

#### D. Transient expression analysis by GUS assay

1. After 3-days co-cultivation, rinse seedlings in sterile water.
2. Place seedlings in 50 ml Falcon tubes with Histochemical GUS staining solution.
3. Apply vacuum for 10 min.
4. Incubate overnight at 37 °C.
5. After staining, rinse seedlings in 70% ethanol to remove excessive stain.
6. Keep seedlings in 70% alcohol for distaining of chlorophyll.

### Recipes

1. YEP liquid medium (1L) 10 g Yeast extract 10 g Peptone 5 g NaCl pH 7.0 Autoclave
2. GM medium (1L) 4.43 g Murashige & Skoog Basal Medium with Vitamins 10 g Sucrose 500 mg MES pH 5.7, autoclave
3. MS sold media (1L) 4.43 g Murashige & Skoog Basal Medium with Vitamins 8 g Agar pH 5.7, autoclave
4. Histochemical GUS stain solution 2 mM Potassium ferrocyanide 2 mM Potassium ferricyanide 100 mM Sodium Phosphate Buffer 500 mg X-Gluc (pre dissolve in dimethyl formamide) 0.1% Triton X-100 1 mM EDTA

## Results and Discussion

Transient expression analysis provides a rapid method to study the function of genes. Transient transformation protocols may also be used to develop stable transformation protocols. In this study, we have reported a rapid and efficient method for transient expression in *Cannabis sativa* seedlings using *Agrobacterium tumefaciens*-mediated transformation. The *Agrobacterium tumefaciens* strain EHA105 carrying the pCAMBIA1301 construct was used to transform cannabis seedlings and the GUS assay was used to detect the transgenics.

Hydrogen peroxide (H_2_O_2_) has been used as a disinfectant for seeds for decades [6]. The 1% H_2_O_2_ was used for both seed sterilization and rapid germination step in solution (unpublished protocol). This presents significant advantage over mercuric chloride or bleach that require additional washing of seeds and germination step in MS medium. This is a very rapid germination method as germination occurs in 24h and seedlings development in 72h. After 3-4 days of incubation in 1% H_2_O_2_ solution, seedlings leave seed coats with two fully opened cotyledons and two immature true leaves (unpublished protocol; Fig. 1); seedlings at this developmental stage were used for transformation. Literature survey displayed that different cultivars of cannabis showed different germination response and revealed optimal germination within 4-7 days by using various germination methods [7] and develop a seedling in 5-15 days or even more. Similarly, Çavusoglu and Kabar [8] showed that exogenous application of H_2_O_2_ to seeds of different plant species increases seed germination rates, coleoptile emergence percentages, radicle and coleoptile elongation, and fresh weights of the seedlings.

The overall workflow for the transient transformation is presented in Figure 2. We have used the intact seedlings (two cotyledons stage and two cotyledons with young true leaves stage) for transformation. To enhance the efficiency, we have used the vacuum infiltration followed by 3-days co-cultivation on MS media. Vacuum infiltration has been shown to enhance the transformation efficiency of *Artemisia annua* seedlings [9]. To detect the gene transformation in cotyledons and true leaves, the GUS activity assay has been done (Fig. 3). GUS analysis revealed that both cotyledons and young true leaves are amenable to transformation. We have carried out the transformation experiment four times. In one independent experiment, we have used around 30 seedlings and out of 30 seedlings at least 20 seedlings showed the GUS activity spots. Previously, Feeney and Punja [10] successfully demonstrated transient transformation of *Cannabis sativa* cell suspension cultures with *A. tumefaciens* strain EHA101 carrying the binary vector pNOV3635 with a gene encoding phosphomannose isomerase, although they failed to regenerate fully transgenic cannabis plants. Wahby et al. [4] reported that hypocotyls tissues were most susceptible to *A. rhizogenes* infection, while young leaves and cotyledons did not, even when the bacteria were stimulated with acetosyringone. These contradicting results may be due to different *Agrobacterium* strains or different cannabis cultivars used in studies.

**Figure 2:**
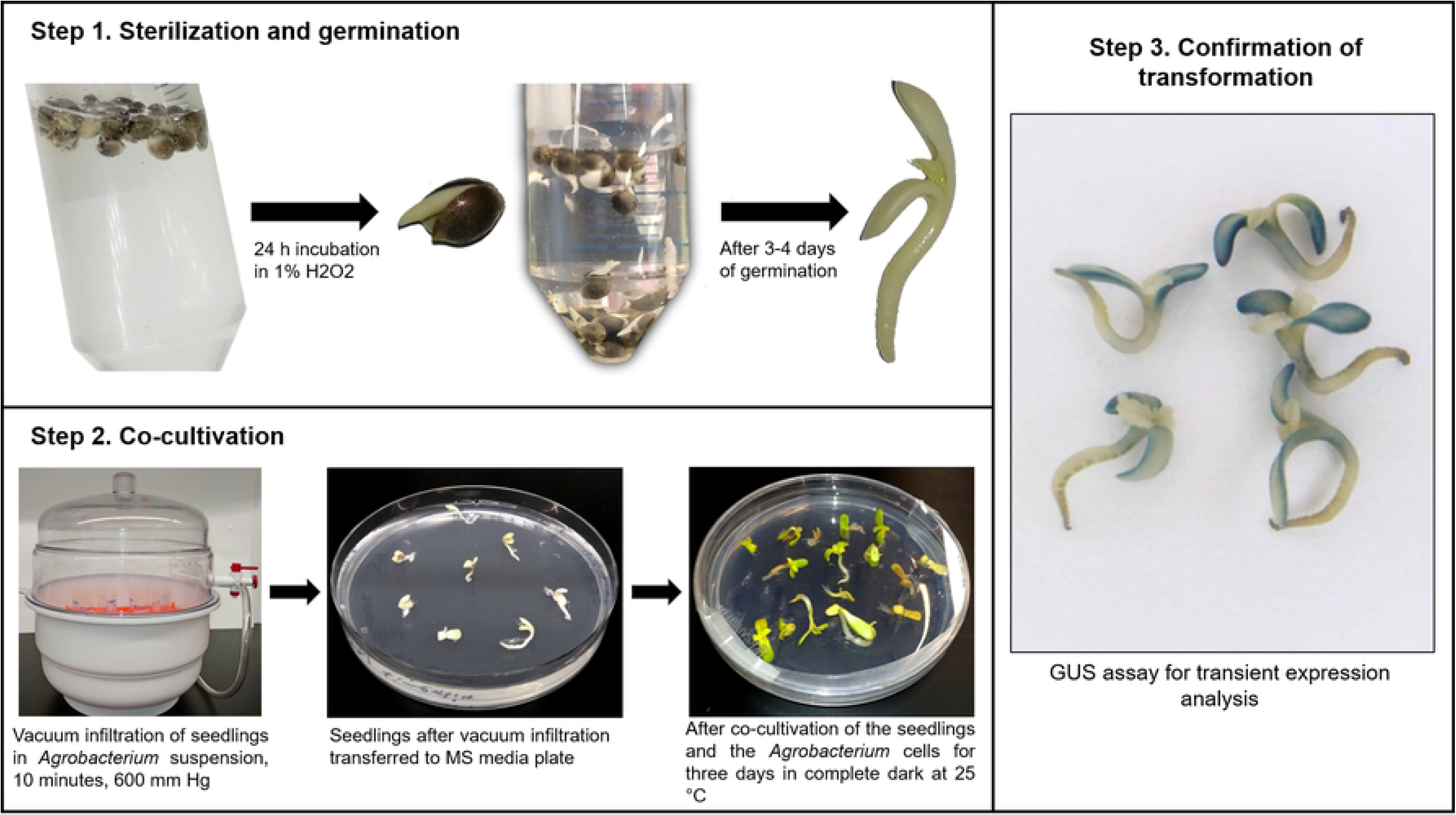
Workflow for *Agrobacterium*-mediated transient transformation of cannabis seedlings. Step 1. Sterilization and germination, seeds are soaked in 1% H_2_O_2_ solution for 24 hours until germination and then transferred into fresh solution. Seeds are then incubated in 1% H_2_O_2_ until both cotyledons and epicotyl are visible. Step 2. Co-cultivation, vacuum applied to seedlings submerged in *Agrobacterium* cell suspension, seedlings are then transferred to MS media plates and incubated for three days in complete dark at 25°C. Step 3. Confirmation of transformation, histochemical GUS assay using transformed seedlings.

**Figure 3:**
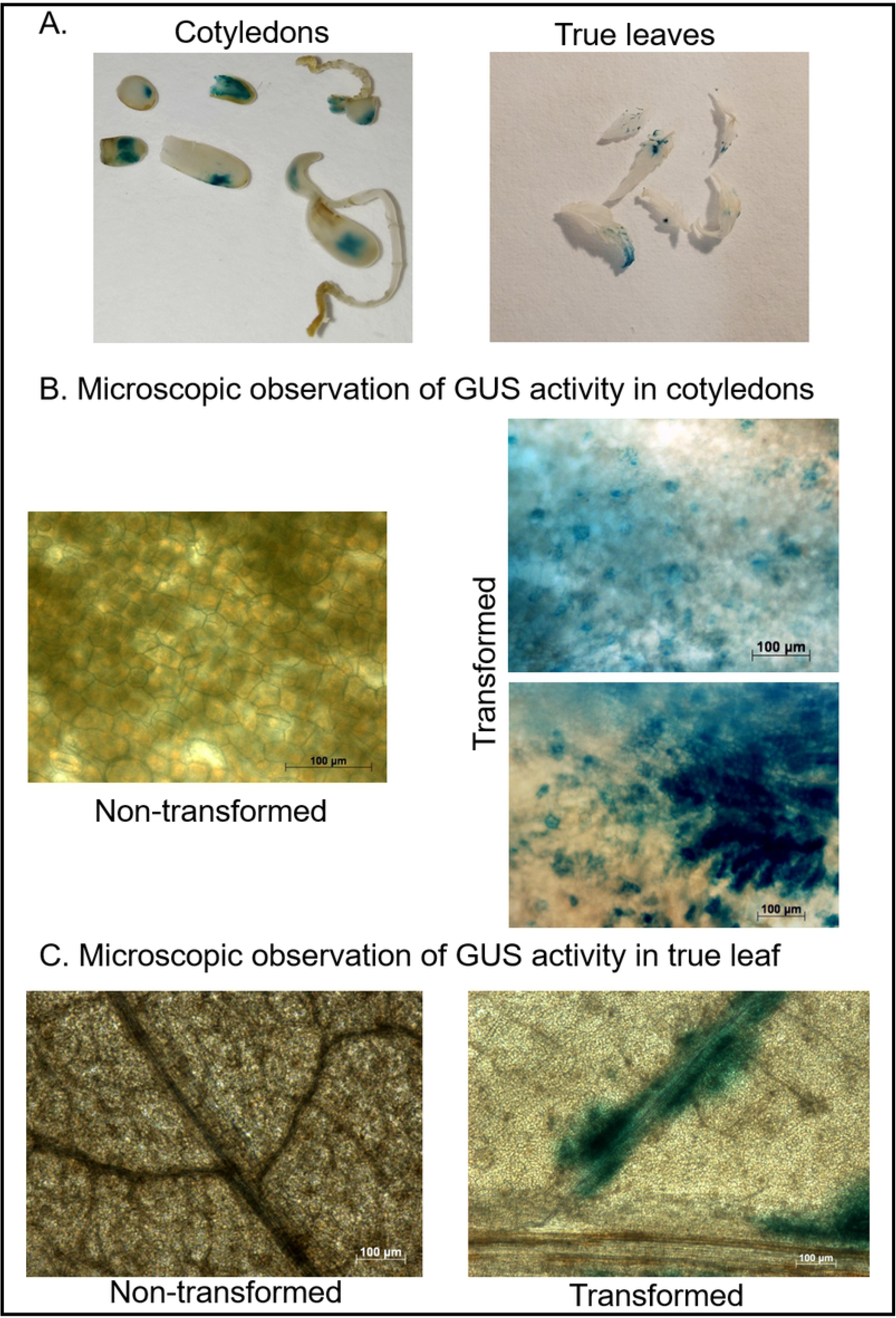
Representative images of GUS activity analysis in cotyledons and leaves tissues of cannabis seedlings to confirm the transient transformation. (A) GUS activity analysis in cotyledons (left panel) and true leaves (right panel). (B) Microscopic observation of GUS activity in cotyledons, non-transformed tissue (left panel) and transformed tissue (right panel). Scale bar 100 μM. (C) Microscopic observation of GUS activity in true leaf, non-transformed tissue (left panel) and transformed tissue (right panel). Scale bar 100 μM.

Comparative qualitative analysis revealed that cannabis seedlings showed less GUS activity than *Nicotiana benthamiana* which suggests that cannabis is less susceptible to *Agrobacterium* infection than *Nicotiana benthamiana* (Fig. 4A). The susceptibility to *Agrobacterium* infection also varied with the different cannabis cultivars. The Nightingale cultivar showed higher susceptibility than the Green Crack CBD and Holy Grail × CD-1 (Fig. 4B). Previously, Feeney and Punja [11] reported that cannabis is amenable to genetic transformation using *Agrobacterium* however the plant is recalcitrant to regeneration, impeding the recovery of transgenic cannabis plants.

**Figure 4:**
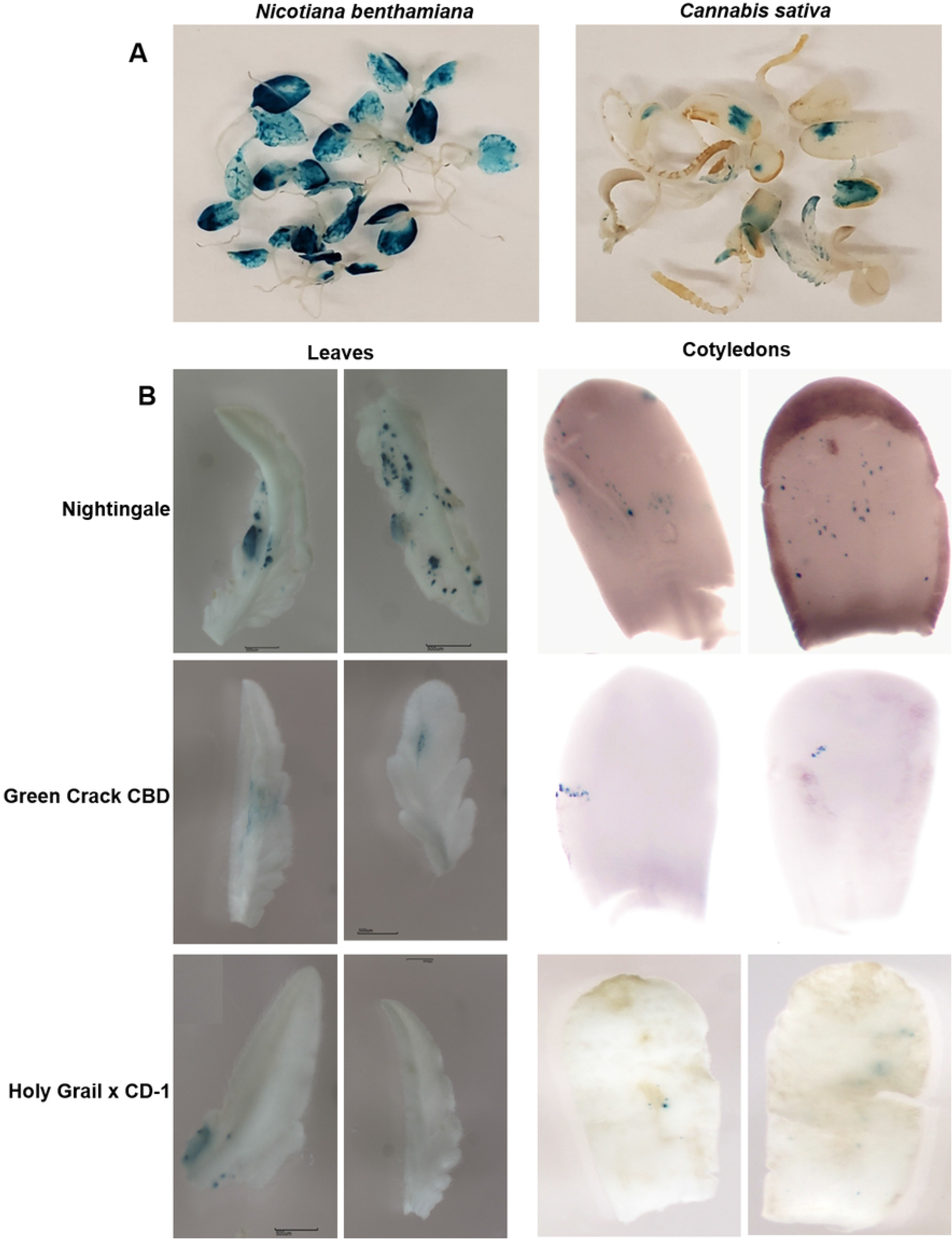
Representative images of comparative transient expression analysis. (A) Comparative transient expression analysis between cannabis and tobacco. (B) Comparative transient expression analysis between different cannabis cultivars.

In conclusion, we have developed an efficient method for transient expression analysis which has potential to be an important tool for gene-function studies and genetic improvement in *C. sativa*.

## Acknowledgements

We thank NSERC and MITACS for funding our work.

## Competing interests

The authors declare that they have no competing interests.

